# A Unified Framework for Batch Correction and Imputation in Modern Mass Spectrometry Proteomics

**DOI:** 10.64898/2026.05.19.726178

**Authors:** Ali Mostafa Anwar, Salma Bayoumi, Leo Lahti, Eleanor Coffey

**Affiliations:** Turku Bioscience Centre, University of Turku and Åbo Akademi University, Turku 20520, Finland; Department of Computing, Faculty of Technology, University of Turku, Turku 20014, Finland

**Keywords:** Batch correction, Proteomics, Mass spectrometry, Non-negative matrix factorization, Missing value imputation

## Abstract

Mass spectrometry-based proteomics, including single-cell applications, are often confounded by batch effects, inter-laboratory variation, and signal drift that can obscure biological interpretation. NMFBatch addresses these sources of unwanted variation through a unified framework that combines non-negative matrix factorization with generalized additive modelling to jointly capture discrete and continuous technical effects from protein expression data. It handles missing values directly through masking and on-the-fly imputation. Benchmarking across reference datasets, plasma proteomics cohorts, simulations, and single-cell data demonstrated that NMFBatch reduced technical variation while preserving biological structure. These results establish NMFBatch as a robust framework for batch correction, harmonization, and integrative proteomics analysis from protein expression data.

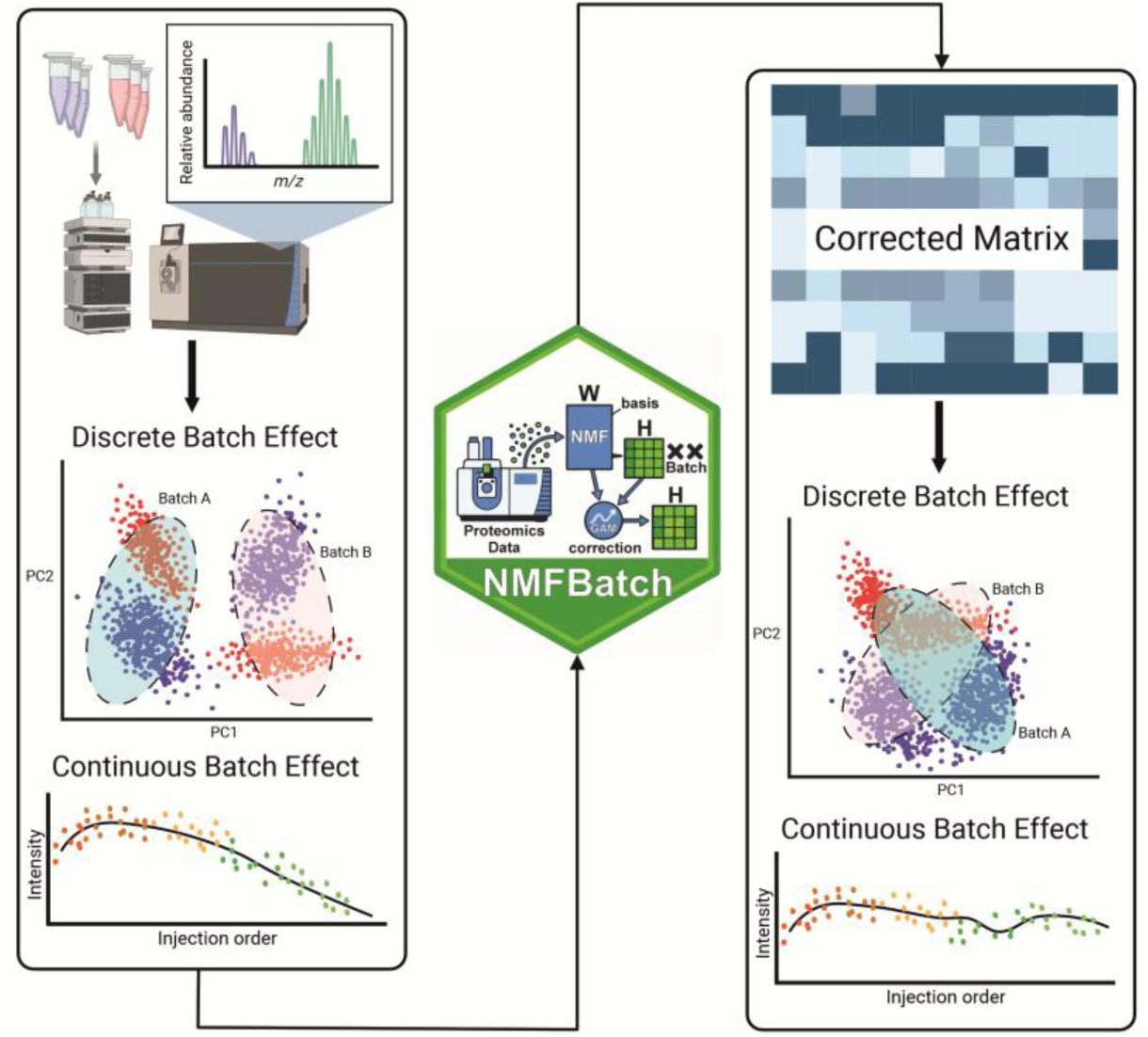

## Introduction

Mass spectrometry (MS)-based proteomics enables the analysis of hundreds of samples and the quantification of thousands of proteins, extending even to single-cell resolution ^1–4^. These developments have expanded the utility of proteomics for the investigation of biological systems and disease mechanisms and placed it at the forefront of technologies for precision medicine ^2,5^. However, the increasing scale and complexity of proteomics studies magnify the impact of technical variability. Quantitative measurements can be affected by differences in sample handling, reagent lots, instrument performance, run order, and inter-laboratory workflows ^6–9^. Collectively, such sources of technical variation are referred to as batch effects. If not adequately controlled, batch effects can distort protein abundance estimates, mask genuine biological differences, reduce statistical power, and limit the reproducibility of proteomics analyses ^7^.

Batch correction in MS-based proteomics is particularly challenging because the data have properties that differ from those of many other high-dimensional omics assays. Large-scale proteomics data typically exhibit two distinct forms of batch effects: discrete batch effects, which manifest as systematic shifts between batches, and continuous batch effects, including progressive signal drift during data acquisition ^9^. These characteristics highlight the need for correction strategies that are specifically suited to the structure of proteomics data. Moreover, missing values are often not missing completely at random (MCAR), rather they are linked to batch structure and other technical factors, known as missing at random (MAR) and may also be missing not at random (MNAR).

Widely used methods based on mean or median adjustment, including ComBat ^10^, often require exclusion of proteins absent in one or more batches, resulting in considerable data loss. To mitigate this problem, missing values are often imputed before correction, which can further distort batch-associated structure ^9,11,12^. Methods such as HarmonizR ^13^ attempt to address this limitation by partitioning the data matrix into subsets based on missingness patterns and subsequently applying ComBat or limma (removeBatchEffect method) ^14^. This reduces the need for pre-correction imputation and minimizes information loss. Nevertheless, these methods remain limited in handling non-linear continuous batch effects. Recognizing this limitation, Čuklina et al. proposed the use of Locally Estimated Scatterplot Smoothing (LOESS) within each batch to model MS signal drift before applying discrete batch correction with ComBat ^9^. This approach was subsequently implemented in the proBatch package.

In this study, we present NMFBatch, a unified framework designed to jointly model discrete and continuous (linear and nonlinear) sources of unwanted variation in large-scale bulk and single-cell MS proteomics experiments directly on the protein level data. NMFBatch handles missing values by masking them during the correction procedure, thereby minimizing feature loss. The framework additionally supports both on-the-fly imputation during batch correction and post-correction imputation without amplifying batch-associated bias. To evaluate performance, we benchmarked NMFBatch against commonly used batch correction approaches ^8^, including ComBat, Median centering, Ratio normalization, RUV-III-C, Harmony, and WaveICA2.0, across multi-laboratory benchmarking datasets ^15,16^ as well as a large-scale real-world dataset. The stability of the method further evaluated using Clinical Proteomic Technology Assessment for Cancer (CPTAC) consortium interlaboratory spike-in datasets ^17^. Finally, we present a case study demonstrating the application of NMFBatch with a real single-cell proteomics dataset ^4^.

## Methods

### NMFBatch Pipeline

The NMFBatch method integrates NMF with flexible GAMs based adjustment for proteomics datasets. The pipeline is applied to proteins data matrices accompanied by sample-level metadata describing batch structure and biological conditions (if available). A full design formula incorporating both biological and batch effects (e.g., ∼ Biology + Batch) or batch only, along with an optional reduced model excluding batch (∼ Biology), was specified to guide downstream adjustment. The workflow begins by estimating the optimal factorization rank to balance model complexity and reconstruction fidelity. Prior to model fitting, proteins exhibiting substantial missingness are removed when such values would prevent reliable NMF. The filtered dataset is then decomposed using NMF into a basis matrix (W) and a coefficient matrix (H). Batch-associated structure in the H matrix is quantified by GAMs, contrasting the full and reduced models, enabling estimation of unwanted variation attributable to batch, which then removed from H, yielding an adjusted latent representation. Finally, the corrected H matrix is used together with W to reconstruct a batch-corrected proteomics expression matrix.

NMFBatch is applied to protein-level data as indicated. Standard normalization procedures, such as reference-channel scaling or median normalization on the peptide level before aggregated to protein level, are compatible. We do not recommend applying additional between-batch normalization after NMFBatch correction, as stacking multiple between-batch normalization procedures may compound distortions and potentially attenuate biologically relevant signal.

Throughout this manuscript, “feature” refers to a quantitative entity represented by a row in the input matrix. When NMFBatch is applied to protein-level data, a feature therefore corresponds to a protein or protein group. The term “feature” does not refer to the latent components of the NMF model, which are referred to as latent factors (or ranks), nor does it specifically refer to precursor or fragment measurements that are subsequently summarized at the protein level.

### NMFBatch Mathematical framework

NMF is a dimensionality reduction technique that decomposes a high-dimensional data matrix into a set of lower-dimensional, non-negative components ^18^. For a given matrix A of size *n × m* (e.g., genes or proteins by samples), NMF approximates A as the product of two matrices, W and H.

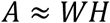

The matrix W has dimensions *n × k*, and H has dimensions *k × m*, where k represents the number of latent patterns (also known as the factorization rank). A defining feature of NMF is the non-negativity constraint imposed on both W and H. This non-negativity leads to a natural additive interpretation of the data: each column of W can be viewed as a “meta-gene” or biological signature, while each row of H reflects how strongly that signature is expressed across the samples _19–21._

In the NMFBatch framework, non-negative matrix factorization is carried out using the NNLM R package ^22^. The underlying optimization approach is alternating non-negative least squares (ANLS), scheme, in which the factorization problem is decomposed into a sequence of non-negative least squares (NNLS) subproblems. By default, NNLM uses a sequential coordinate-wise descent (SCD) algorithm, which updates one element of the factor matrices at a time while keeping the others fixed, repeating this process until convergence. To mitigate overfitting, NNLM incorporates multiple regularization strategies applicable to both W and H. L2 (ridge) regularization, which controls magnitude and smoothness; angular regularization, which reduces correlations among columns; and L1 (lasso), which promotes sparsity in the matrices. The resulting optimization problem can be written as:

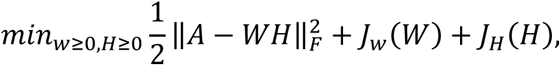

where *A* ∈ ℝ*^n^*^×*m*^, *W* ∈ ℝ*^n^*^×*k*^, *H* ∈ ℝ*^k^*^×*m*^, and *J_w_*, *J_H_*(*H*), are regularization penalties as defined in _22._

An important feature of NNLM is its ability to handle missing data. If the input matrix A contains missing values (NAs), these entries are ignored (masked) during error computation and parameter updates. After factorization, the missing entries can be estimated using the reconstructed matrix (W × H), imputing values based on learned patterns (on-the-fly imputation).

### Generalized Additive Models (GAMs) Based Adjustment of Latent NMF Components

We corrected batch effects at the level of latent NMF components by operating on the coefficient matrix *H*. Each component vector ℎ*_k_* (across samples) modeled independently using generalized additive models (GAMs) with penalized regression spline and smoothing parameter estimation via restricted maximum likelihood (REML).

For each component *k,* we modeled the expected value as:

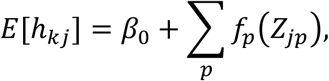

where *Z_jp_* denotes the *p*-th covariate for sample j, and *f_p_*(·) denote model terms, which may be specified as smooth functions (e.g., spline-based terms) or parametric linear effects depending on the model formula. This defines the full model:

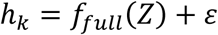

When specified, a reduced model excluding selected covariates was also fitted:

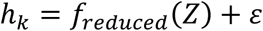

The estimated batch contribution was defined as the difference in fitted values from the GAMs defined as:

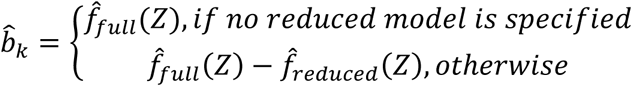

Finally, batch corrected component values could be obtained as:

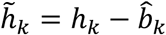

where, the final corrected coefficient matrix can be noted as *H̃*.

After batch effect correction in the latent space, the adjusted coefficient matrix *H̃* are used for data reconstruction while keeping the original basis matrix W fixed. The corrected latent matrix is allowed to be unconstrained in ℝ, yielding a semi-NMF formulation in which the reconstructed data defined as *A*^∗^ = *WH̃*.

### Rank Selection in NMFBatch

To determine the optimal factorization rank (k), we performed a rank-selection procedure across a predefined range of candidate ranks (default: k = 2–10). For each rank (k), after matrix factorization, the reconstructed matrix was generated from W and H, and missing values were restored to their original locations. Reconstruction performance for each rank is quantified using the relative residual error derived from squared differences between observed and reconstructed matrices. Total explained variance is computed as one minus the normalized reconstruction error.

When sample metadata are available, latent-factor diagnostics are computed to assess preservation of biological structure and separation of batch-associated variation within the latent representation. Biological variance is estimated within batches and averaged across batches to quantify retained biological structure, while batch variance is estimated within biological conditions and averaged across conditions to evaluate remaining batch-driven structure across ranks.

Factor-level summary metrics are also calculated. Factor purity quantifies the separation between batch-associated and condition-associated contributions across latent factors. Factor coverage quantifies the strength of the dominant structured signal within each latent factor, independent of whether the signal originates from batch or biological condition.

For each candidate rank, diagnostic plots are generated for visualization of model behavior. In this study, rank selection follows criteria prioritizing high explained variance, low reconstruction error, and stable factor purity and factor coverage across both biological and/or batch-associated structure.^23^

### NMFBatch R package implementation

NMFBatch is implemented as an openly licensed R package and available on Codeberg (https://codeberg.org/AliYoussef/NMFBatch). The package supports the SummarizedExperiment data structure ^24^, improving compatibility with many Bioconductor-based tools. Additionally, the package provides functionality for generating batch effect– focused diagnostic plots, including PCA plots, heatmaps, and boxplots. Comprehensive documentation covering all NMFBatch functions and the full workflow is available as reproducible vignettes.

### Benchmarking Datasets and Evaluation Metrics

#### Quartet multi-lab datasets

We used six preprocessed datasets derived from Quartet reference materials, as multi-laboratory benchmarking datasets ^15,16^ and were adopted directly from Chen et al. ^8^ and downloaded from Figshare database ^25^. This dataset contains three technical replicates of four biological conditions of immortalized B-lymphoblastoid cell lines (LCLs), derived from a quartet family of parents and monozygotic twin daughters (F7, M8, D5, and D6). In this study, we used preprocessed MS datasets generated in earlier work ^8,15,16^, where raw files were processed using MaxQuant (v2.1.3.0) with default parameters and searched against the human UniProt database (release 2022-08-08).

Datasets DAT1-3 were generated using data-dependent acquisition (DDA) strategy and analyzed directly. In contrast, DAT4-6 was produced using a DDA-library-based data-independent acquisition (DIA) workflow, and the resulting DIA MS files were analyzed using MaxDIA implemented within the MaxQuant platform. To model a balanced experimental design, the four sample types (D5, D6, F7, and M8) were distributed in equal proportions across all batches. In the confounded scenario, however, each batch included only two sample types (D6 and D5 or F7 or M8), resulting in batchs and samples that are partially confounded ^8^. Prior to batch-effect correction, the protein-level was aggregated from peptide level data using MaxLFQ, then preprocessed using a log transformation followed by per-sample normalization of total intensities, without missing-value imputation as described in the original study ^8^.

#### ChiHOPE dataset

As a real-case application, we reused the dataset processed in the same study by Chen et al. ^8^ and downloaded from Figshare database ^26^ which analyzed the Chiglitazar-perturbed Human multi-Omics ProfilE (ChiHOPE) cohort. The dataset comprises plasma proteomics profiles from 750 individuals with type 2 diabetes (T2D), including 274 females and 476 males, aged between 23 and 70 years. Three categories of quality control (QC) samples were injected in each batch together with the T2D samples. The first QC group (PM) consisted of pooled plasma samples generated from the study cohort. The other two QC groups (P10 and P11) were plasma samples derived from healthy male and female donors, respectively.

Samples were measured in DIA mode using an EASY-nLC 1200 system coupled to a Q Exactive HF-X. In each 96-well plate, study samples were co-injected with two PM samples, one P10 sample, and one P11 sample, yielding 1,495 MS runs overall ^8^. The resulting DIA datasets were analyzed using DIA-NN (v1.7.0)^27^ and the UniProt human protein database (release dated 17 December 2019). In this dataset, the corrections for all methods done with all samples included (study samples and QC samples), however the benchmark analysis (Evaluation sub-section) are done only on the QC samples. Prior to batch-effect correction, the protein-level was aggregated from peptide level data using MaxLFQ, then preprocessed by log transformation followed by per-sample normalization of total intensities as described in the original study ^8^.

#### Assessment of NMFBatch in the CPTAC Proteomic Cancer Spike-In Protein Dataset

To assess the impact of NMFBatch on **ground-truth** spike-in proteins, we used data from the interlaboratory CPTAC consortium study ^17^ bulk label-free proteomics spike-in data. Forty-eight human proteins were spiked at five different concentrations, ranging from 0.25 *fmol/µL* to 20 *fmol/µL*. For each concentration, three replicate samples were distributed to three different laboratories, which capture interlaboratory variability. The peptide dataset was sourced from https://github.com/statOmics/GMFProteomicsPaper ^28^. Peptides with more than 90% missing values and protein groups supported by fewer than two peptides were removed. We used Huber’s robust M-estimator to aggregate peptide-level intensity values into protein-level abundances. We then retained proteins with a missing value proportion of ≤30%.

#### Leduc pSCoPE Single cell proteomics dataset

As a single-cell proteomics data, we utilized a dataset from Leduc et al. ^4^. The data were generated using pSCoPE technology with TMT labeling and data-dependent acquisition performed on a Thermo Scientific Q Exactive instrument. The dataset, downloaded through the scpdata package ^29^, contained quantitative measurements for 2844 proteins across 1543 single cells. Protein intensities were first normalized to the reference channel and log2-transformed ^29^. Because the dataset had already undergone normalization and contained negative values, the minimum absolute value in the dataset, together with a small offset (1 × 10⁻⁶), was added to all entries to ensure compatibility with NMFBatch. Proteins with missing values greater than or equal to 70% were subsequently removed. This resulted in the removal of 1113 of 2844 proteins (39%). Liquid chromatography (LC) batch information and TMT labels were incorporated into the full NMFBatch model, with no reduced model specified.

### Batch-effect correction Methods

We systematically benchmarked NMFBatch against six established batch-effect correction algorithms, widely used proteomics (ComBat, RUV-III-C, Harmony, Ratio normalization, median centering, and WaveICA 2.0). For the ChiHOPE dataset, we additionally evaluated all of the methods in combination with per-batch LOESS correction. ComBat utilizes an empirical Bayes framework to model and remove additive and multiplicative batch effects while preserving biological differences ^10^. RUV-III-C removes unwanted variation by estimating latent factors from sample labels and negative-control features, which are subsequently regressed out ^6^. For RUV-III-C in the Quartet dataset, negative-control features were defined via two-way ANOVA, selecting protein groups that lacked significant biological variation across samples or exhibited batch-driven variation without consistent sample-level effects. Harmony corrects batch effects by embedding data in a shared low-dimensional space and iteratively aligning samples to minimize technical variation while preserving biological structure ^30^. Although originally developed for single-cell RNA-seq data, it is applicable to other omics datasets. Ratio normalization was performed by subtracting the mean expression of a group of samples within each batch from all other samples.

Median centering applies a location shift to align per-feature medians across batches to a common reference. WaveICA 2.0 uses wavelet-based decomposition to separate technical and biological signals and can remove both inter-and intra-batch effects without requiring explicit batch labels ^31^.

#### Principal Variance Component Analysis (PVCA)

To quantify the relative contributions of technical and biological sources of variation, we applied a PVCA framework. Prior to analysis, we excluded features exhibiting zero variance or containing missing values and performed principal component analysis (PCA) on centered and scaled data. We then selected a subset of the principal components to capture at least 60% of the total variance, with a minimum inclusion of five components to ensure stability of variance estimates. For each component, we used linear mixed-effects models to partition the variance with batch and sample type specified as random effects, including their interaction. Finally, we normalized variance components to the total variance and aggregated across components using weights proportional to the variance explained. In this PVCA we did not interpret the interaction term between batch and sample type separately and was instead included in the residual variance, resulting in three reported components: batch, biological (sample), and residual variance.

#### Silhouette analysis

We then used Silhouette analysis to assess the strength of cluster structure defined by categorical labels (batch and condition). Using a subset of reduced-dimensional space using PCA capture at least 60% of the total variance, with a minimum inclusion of five components. Each observation receives a Silhouette width based on the difference between its average dissimilarity to observations within the same group and to those in the nearest alternative group. This provides a measure of cluster membership quality relative to competing clusters. We summarized the overall clustering strength for a grouping variable as the mean Silhouette width across observations, where higher values indicate stronger separation and lower values indicate greater overlap.

#### Intraclass Correlation Coefficient (ICC)

To evaluate the batch effect on feature-level reproducibility, we used the ICC. For each feature, we partitioned variability using a one-way random-effects analysis of variance model with sample type as the grouping variable. After, we estimated between-group and within-group variance components using mean squares derived from replicate observations. Finally, we defined the ICC as the proportion of total variance attributable to between-group variability, providing a measure of reproducibility across biological groups. ICC values were constrained to the interval [0, 1] to ensure interpretability, therefore, higher median ICC (closer to 1) indicates that biological sample membership explains a larger fraction of total variance, reflecting better within-sample reproducibility after correction.

For the benchmark evaluation analysis, we removed proteins with missing values or zero standard deviation from all datasets produced from batch correction methods, and we only retained proteins common to all datasets.

## Results

### Experimental design overview

We evaluated NMFBatch, a latent-space method that combines non-negative matrix factorization (NMF) with generalized additive models (GAMs), for correction of technical variation in proteomics data (Fig. 1A). The evaluation covered discrete batch effects, continuous signal drift across injection order, missing values, sparse single-cell data, computational stability, and the risks of overcorrection and overfitting.

**Figure 1:**
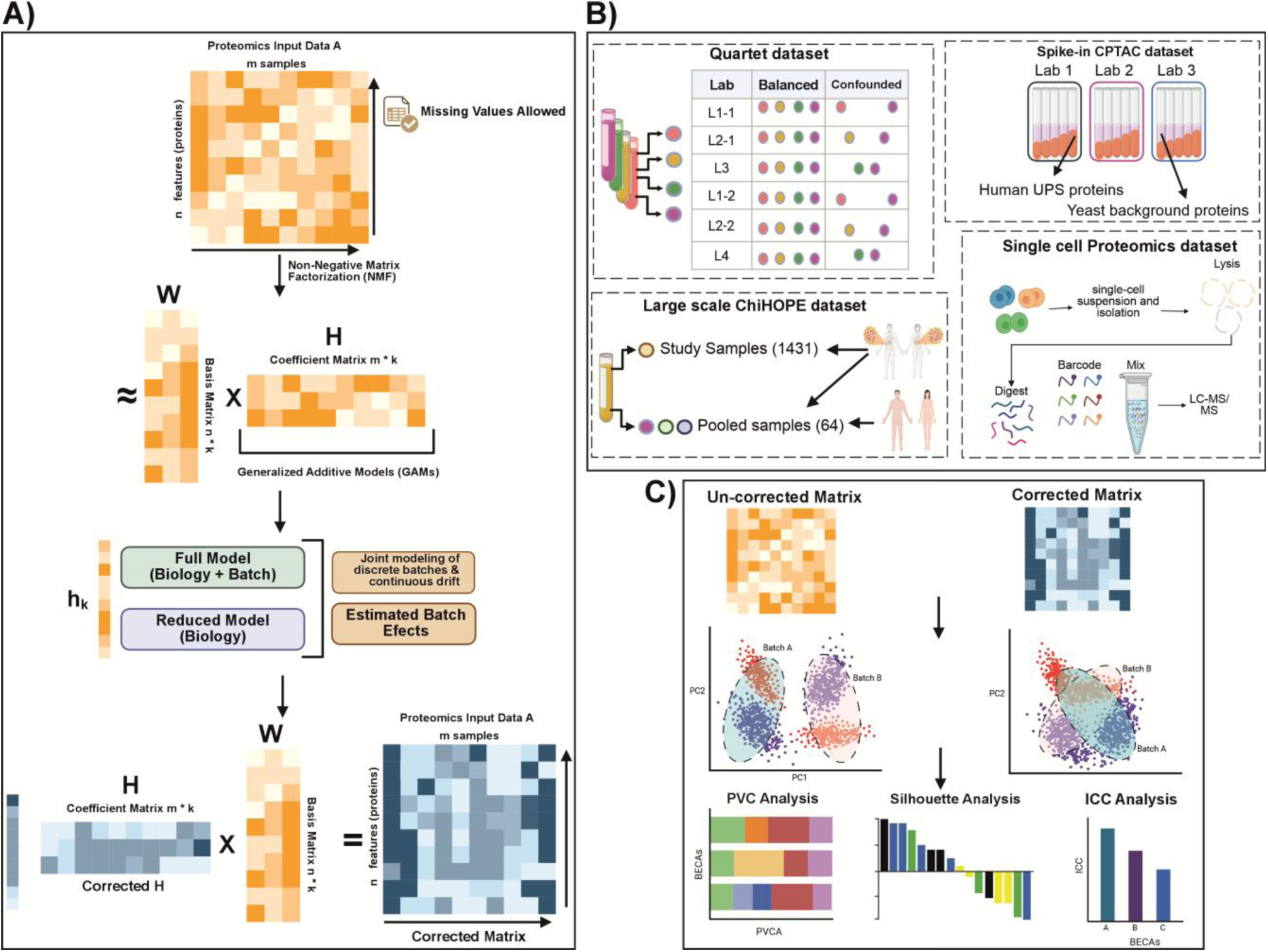
NMFBatch Framework and Overview of the Study Design Schematic overview of the NMFBatch workflow. Starting from a proteomics dataset and sample metadata containing biological and batch variables, the data undergo filtering and rank estimation. NMFBatch then performs NMF to decompose the matrix into latent factors W and H. Batch effects are estimated in H using GAMs with specified design formulas, and the corrected H is recombined with W to reconstruct the batch-corrected proteomics matrix. B) Four datasets were used for benchmarking. The Quartet Project dataset includes balanced (72 samples) and confounded (36 samples) designs. The ChiHOPE plasma proteomics cohort includes 750 individuals and 1,495 mass spectrometry runs. The CPTAC interlaboratory spike-in dataset contains 48 human proteins at five known concentrations measured across three laboratories. The pSCoPE dataset from Leduc et al. ^4^ contains quantitative measurements of 2,844 proteins in 1,543 single cells from three cell populations. C) Evaluation metrics. Batch-correction performance was assessed at both the sample and feature levels using PVCA, silhouette analysis in PCA space accounting for at least 60% of total variance, and ICC. Created in BioRender. Youssef, A. (2026) https://BioRender.com/7ttotr2 and subsequently modified for layout and formatting in CorelDRAW.

To examine the performance of this method, we benchmarked NMFBatch against six established batch-correction methods ^8^, ComBat, median centering, ratio normalization, RUV-III-C, Harmony and WaveICA 2.0, across two datasets. We used the Quartet multi-laboratory reference dataset ^8,15^ in two settings: a balanced design in which all four sample types were represented equally across six batches, and a confounded design, in which each batch contained only two sample types. We also used the ChiHOPE plasma proteomics cohort (8), which included 750 individuals and 1,495 mass spectrometry runs across three batches. Together, these datasets represent controlled multi-laboratory and large-scale cohort settings (Fig. 1B).

We assessed the batch correction methods performance on sample-level using principal variance component analysis (PVCA) and silhouette analysis in principal component analysis (PCA) space. The selected principal components explained at least 60% of the total variance. Next, we assessed the methods on protein-level using the intraclass correlation coefficient (ICC) and visually examined the data using PCA, UMAP, and clustered heatmaps.

To test whether NMFBatch preserved known biological signals without overcorrecting, we analyzed the CPTAC interlaboratory spike-in dataset ^17^. For each protein, we compared the proportions of variance associated with spike-in concentration and laboratory before and after correction. Moreover, we performed repeated-initialization and permutation analyses to assess computational stability and potential overfitting (Fig. 1B and C).

Finally, we applied NMFBatch to the pSCoPE single-cell proteomics dataset from Leduc et al. ^4^. This dataset contained quantitative measurements for 2,844 proteins across 1,543 single cells acquired across two LC batches, 1543 TMT labelling sets (Fig. 1B). It provided a test of performance under substantial sparsity and multiple technical sources of variation, including liquid chromatography batches and TMT labeling sets.

### NMFBatch removes batch-associated variation while preserving biological stracture

We compared NMFBatch with six batch-correction methods in the Quartet and ChiHOPE datasets on the protein level data. Before correction, the Quartet samples clustered strongly by batch in PCA space (Fig. 2A and Fig. S1). PVCA attributed 97% and 97.1% of the total variance to batch effects in the balanced and confounded Quartet datasets, respectively (Fig. 2B, Table S1). Similarly, in the uncorrected ChiHOPE data, batch effects explained for 71.1% of the variance, whereas sample-related biological variation contributed 6.3% (Fig. 2C, S2).

**Figure 2.**
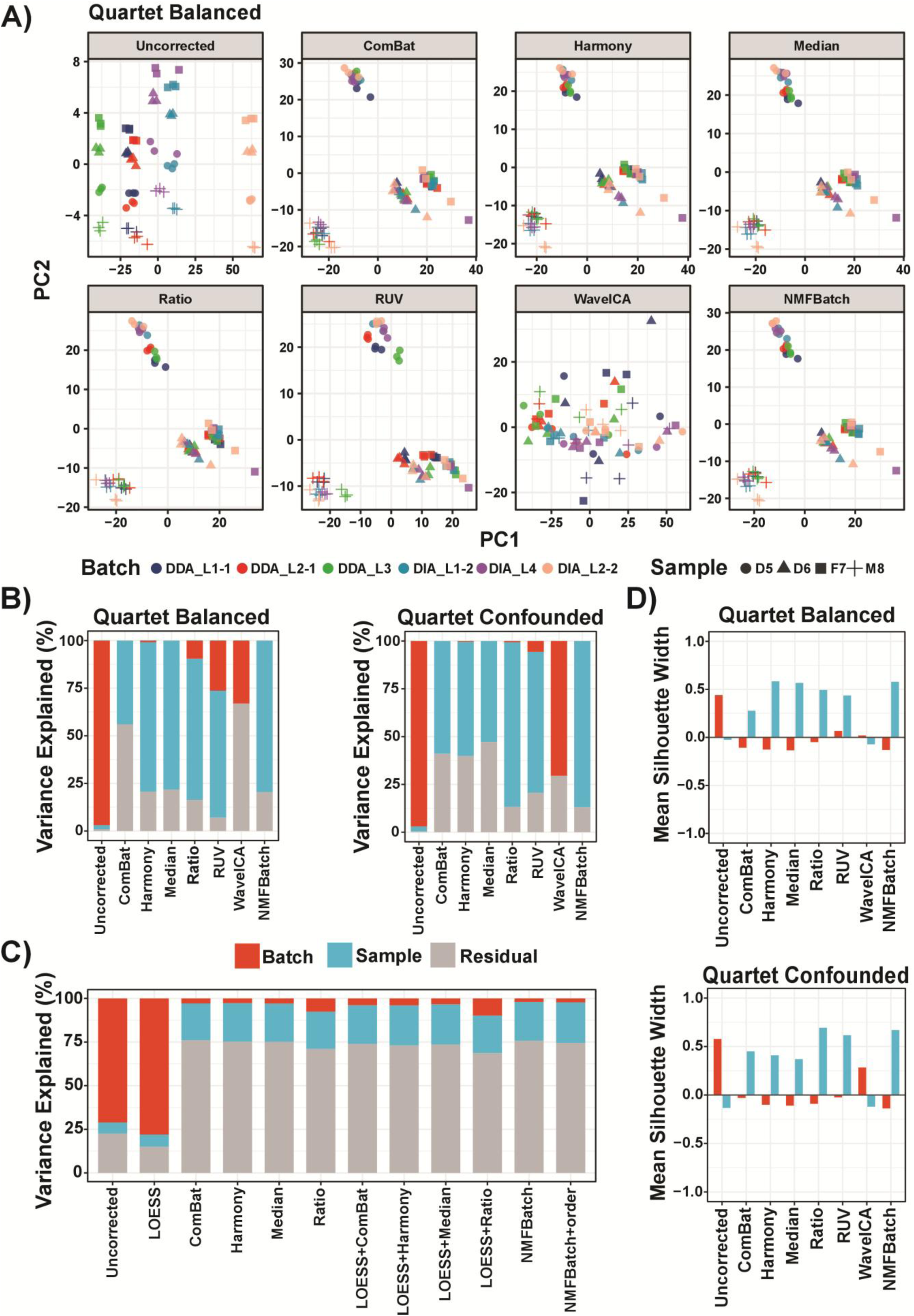
Benchmarking of NMFBatch against established correction methods in balanced, confounded and cohort-scale proteomics datasets. (A) PCA of the balanced Quartet dataset before correction and after correction with ComBat, Harmony, median centering, ratio normalization, RUV-III-C, WaveICA, and NMFBatch. Points are colored by batches. Batches are marked by the acquisition mode data-dependent acquisition and data-independent acquisition (DDA or DIA) and the laboratory number. Points are shaped by sample type (D5, D6, F7, and M8) biological conditions. The uncorrected data show pronounced clustering by batch, whereas correction with NMFBatch reduces batch-driven separation while preserving separation by sample type. B) PVCA of the balanced and confounded Quartet datasets following correction with each method. Bars represent the proportions of variance explained by batch (red), sample (blue) and residual (grey) components across principal components accounting for at least 60% of total variance. NMFBatch reduces batch-associated variance to near zero while preserving the largest or near-largest sample-associated component across both experimental designs. (C) PVCA of the ChiHOPE cohort before correction, after LOESS correction alone, after LOESS followed by alternative discrete batch-correction methods, and after NMFBatch fitted either with batch only or with both batch and injection-order terms. NMFBatch yields low residual batch- associated variance while maintaining competitive recovery of biologically attributable variance. (D) Mean silhouette widths for batch and sample labels in the balanced and confounded Quartet datasets after correction with each method. More negative or near-zero batch silhouette widths indicate improved batch mixing, whereas larger sample silhouette widths indicate better biological separation. Across both designs, NMFBatch achieves low batch silhouette values together with high sample silhouette values.

After NMFBatch correction, samples from different batches mixed more closely in PCA space, while separation by biological sample type remained visible in both Quartet balanced and confounded designs and in ChiHOPE data (Fig. 2A; Fig. S1 and S2.)

PVCA supports the visual results. In the balanced Quartet design, NMFBatch reduced batch-associated variance to approximately zero while recovering the largest biological-associated variance among the tested methods, at 79.6% (Fig. 2B, Table S1). Harmony retained 0.85% batch variance and 78.5% biological variance. Median centering also reduced batch variance to zero and 78.3% restored biological variance.

The difference between methods was larger in the confounded design. NMFBatch reduced batch-associated variance to approximately zero and recovered 87% of the biological-associated variance, the highest value observed among all methods (Fig. 2B, Table S1). Although Harmony reduced batch variance to 0.4%, it recovered only 59.8% of the biological variance. Median centering also completely removed batch variance but recovered biological variance, at 52.8%.

In the ChiHOPE data, when we NMFBatch corrected only the discrete batch effects, it achieved the lowest residual batch-associated variance of any single-step method, reducing it to 2.1%. The corresponding values were 2.8% for Harmony, 2.9% for ComBat, 3.01% for Median centering, and 7.7% for ratio (Fig. 2C and Table S2). NMFBatch also recovered 22.2% of the biological variance.

Silhouette analysis provided an orthogonal assessment of the same clustering pattern, and the results were broadly consistent with those observed in PVCA. In the balanced Quartet design, NMFBatch and Harmony achieved a biological-cluster silhouette score of 0.6, together with negative batch silhouette scores, indicating weak clustering by batch (Fig. 2D, Table S3). Under the confounded design, NMFBatch and ratio normalization yielded the highest sample silhouette scores, at 0.7, whereas Harmony dropped to 0.41. In the ChiHOPE cohort, NMFBatch produced the most negative batch Silhouette score (−0.008), with median centering showing a similarly low value (−0.002), while Harmony and ComBat retained slightly positive batch silhouette scores (Table S4).

Protein-level ICC evaluation results were also consistent with the sample-level analysis. In the balanced Quartet design, NMFBatch achieved a median ICC of 0.50, higher than Median centering (0.49), Harmony (0.49), ratio normalization (0.46), RUV-III-C (0.42), and ComBat (0.15) (Fig. S3, Table S5). In the confounded design of NMFBatch, where the median ICC reached 0.69, compared with 0.64 for ratio normalization, 0.58 for RUV-III-C, 0.39 for Harmony, 0.34 for Median centering, and 0.31 for ComBat (Fig. S3, Table S5).

Overall, NMFBatch consistently reduced batch-associated variation while retaining sample-associated structure. Its advantage was most evident in the confounded Quartet design.

### Joint modelling of batch effects and injection-order drift improves correction in the ChiHOPE dataset

The ChiHOPE data showed nonlinear intensity changes across 1,495 injections, together with discontinuities at batch boundaries (Fig. 3A). The uncorrected data and the data corrected with LOESS alone still showed technical structure in PCA space (Fig. 3B).

**Figure 3.**
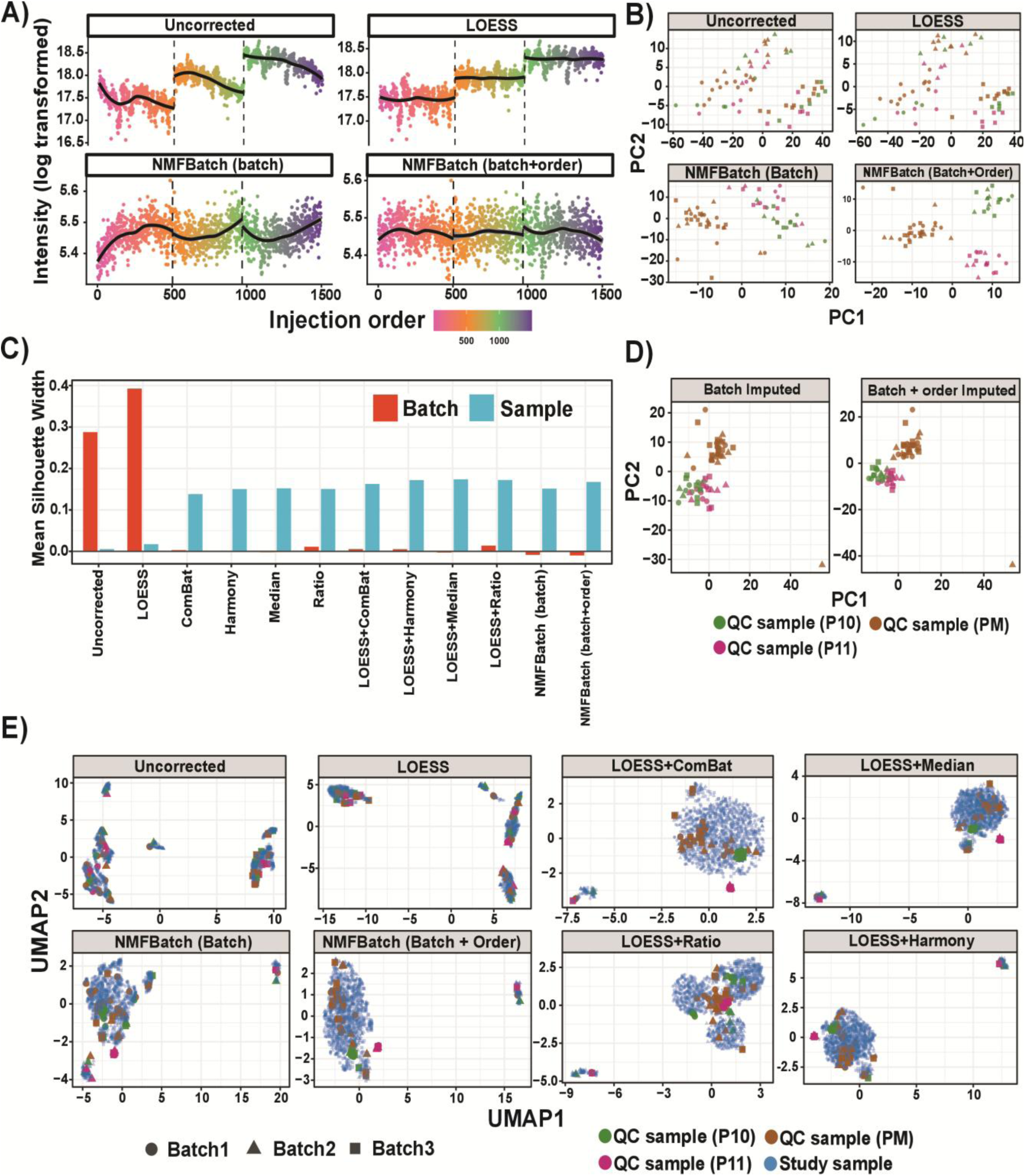
Joint modelling of discrete batch effects and injection-order drift by NMFBatch in the ChiHOPE cohort. (A) Representative protein-intensity trends across injection order in ChiHOPE before correction and after correction using LOESS, NMFBatch with batch effects only, or NMFBatch with both batch and injection-order terms. Colors represent injection order, vertical dashed lines mark batch cleaning instrument transitions, and black smooth curves indicate global intensity trends across the run. (B) PCA of QC samples before correction, after LOESS correction, and after NMFBatch correction with batch-only or batch-plus-order models. Points are colored by QC sample type.

When correction was limited to discrete batch effects, all methods produced moderate separation of QC samples in PCA space (Fig. 3B, Fig. S2). When applying LOESS, it corrects only the MS signal drift effectively, although a small deviation remained between injections 500 and 1000, when combined with other methods (Fig. S4). This deviation was not evident when NMFBAtch modeled batch and injection order jointly (Fig. 3A).

Adding the injection-order term to NMFBatch increased sample-associated variance from 22.2% to 23.2% in the PVCA (Table S2). Silhouette analysis showed that NMFBatch, LOESS+medain centering, LOESS+ratio, LOESS+Harmony produced the highest sample Silhouette score, all approximately 0.17 (Fig. 3C; Table S4). The two NMFBatch models produced the lowest batch silhouette scores. Moreoever, enabling on-the-fly imputation produced QC PCA patterns (Fig. 3D) that were similar to those obtained without imputation (Fig. 3B).

Finally, UMAP analysis of the study and QC samples gave the same general pattern (Fig. 3E). Batch-associated organization remained visible in the uncorrected and LOESS-only data. It was reduced by the two-step correction procedures (LOESS+batch correction method) and by NMFBatch when injection order was modelled with a penalized spline.

These results show that NMFBatch can model discrete batch shifts and continuous injection-order drift in one framework. Because GAM smothers are penalized, and injection-order term can reduce toward zero when the data provide little evidence of drift, limiting unnecessary adjustment.

NMFBatch reduces technical dispersion while retaining separation between QC sample types. (C) Mean Silhouette widths for batch and sample labels in the ChiHOPE dataset after correction with the indicated methods. Lower batch Silhouette values indicate improved batch mixing and higher sample Silhouette values indicate stronger preservation of biological structure. NMFBatch combines low batch Silhouette values with competitive sample Silhouette scores. (D) PCA of matrices imputed during NMFBatch (on-the-fly imputation) correction under the batch-only and batch-plus-order models. The corrected imputed matrices retain separation between QC sample classes. (E) UMAP representation of all ChiHOPE samples before correction and after correction using LOESS alone, LOESS followed by ComBat, median centering, ratio normalization or Harmony, and NMFBatch fitted with batch-only or batch-plus-order models. Points are colored by sample type and shaped by batch. NMFBatch reduces batch-associated structure while preserving coherent organization of the biological sample classes.

### Missing-value masking allows correction without requiring a complete matrix

Many batch-correction methods require a complete or nearly complete matrix. This can require either removal of proteins that are missing in one or more batches or imputation before correction. NMFBatch instead masks missing entries during NMF optimization.

We assessed this function in ChiHOPE dataset by enabling on-the-fly imputation and retaining all proteins. The batch-only model produced 1.68% batch-associated variance and 21.52% sample-associated variance. The batch-plus-order model produced 2.36% batch-associated variance and 22.48% of sample-associated variance. These values were close to those obtained in the filtered-proteome analysis (Table S2). Silhouette analysis of the full proteome likewise yielded negative batch scores (−0.026) and sample scores of 0.14 and 0.16 for the two models (Table S4). In addition, visual inspection indicated effective removal of both MS signal drift and discrete batch effects (Fig. 3D, Fig. S5).

Thus, masking missing values allowed NMFBatch to correct an incomplete protein matrix while retaining proteins that would otherwise have been excluded. In this dataset, the resulting batch and sample metrics were like those obtained after filtering to a complete matrix

### NMFBatch is stable across random initializations and does not show evidence of label-driven overfitting

We used CPTAC interlaboratory spike-in dataset ^17^ to assess whether NMFBatch preserved known concentration-dependent signals without introducing overcorrection including on-the-fly imputation. After correction, the spike-in proteins remain concentrated in the region of high condition-variance, while laboratory-associated variance decreased (Fig. 4A, B).

**Figure 4.**
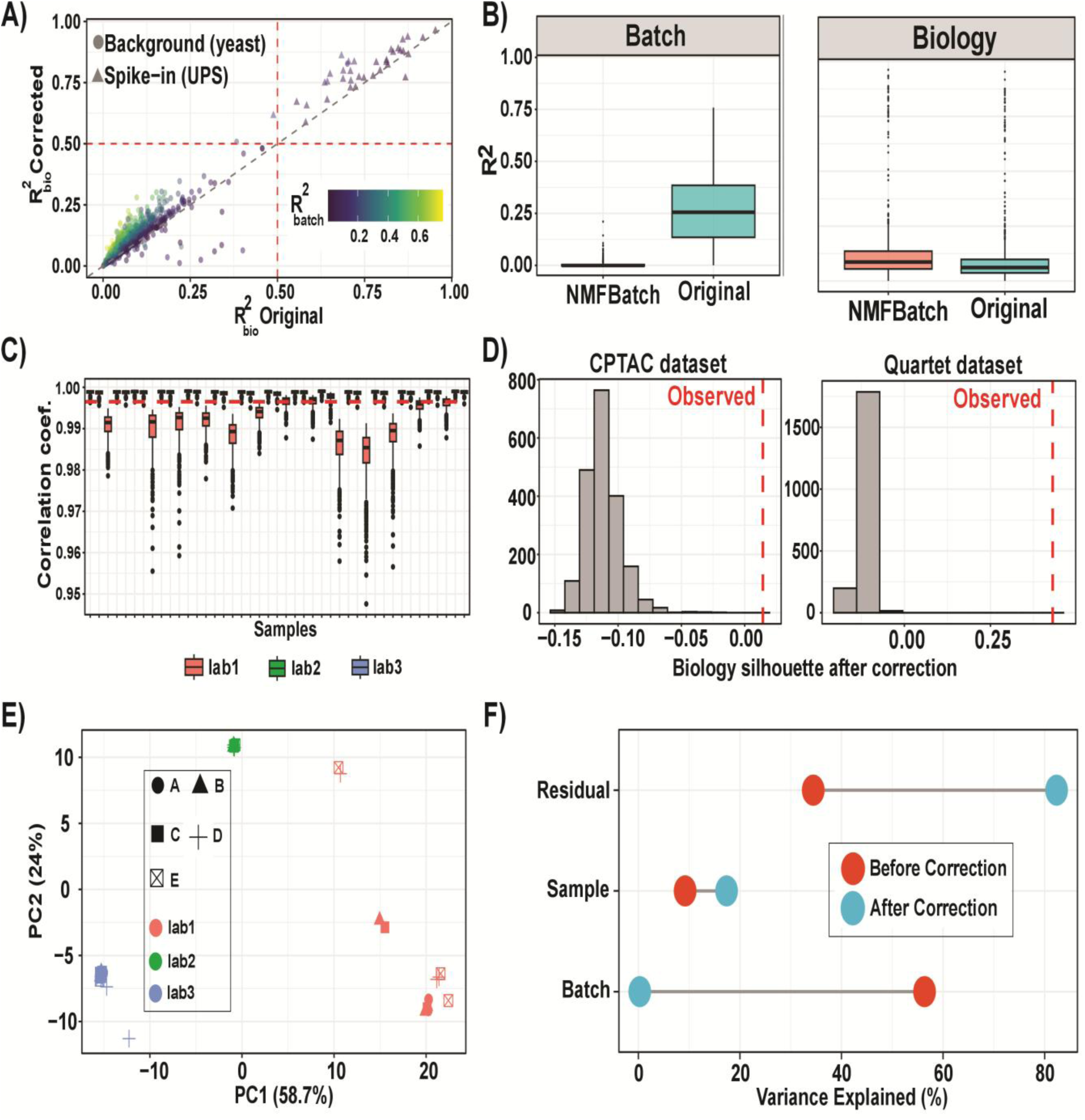
NMFBatch preserves concentration-dependent biological signal and shows stable behaviour across repeated runs (A) Comparison of protein-level condition-associated variance before and after NMFBatch correction in the CPTAC interlaboratory spike-in dataset. Each point represents one protein and is coloured by batch-associated (*R*^2^). Circles indicate background yeast proteins and triangles indicate UPS spike-in proteins. Proteins close to the diagonal retain similar condition-associated variance after correction, indicating preservation of biological concentration-dependent signal. (B) Boxplots showing the distribution of protein-level (*R*^2^) values attributable to batch and biology before and after NMFBatch correction. Batch-associated variance is reduced after correction, whereas biology-associated variance is largely maintained. (C) Distribution of pairwise sample correlation coefficients across 500 independent NMFBatch runs on the CPTAC dataset, stratified by laboratory. Correlations remain tightly clustered near 1, indicating high reproducibility across random initializations. (D) Permutation analysis of biology silhouette scores following NMFBatch correction in the CPTAC (left panel) and Quartet (right panel) datasets. Histograms show the null distributions generated by shuffled labels, and dashed red lines indicate the observed Silhouette scores. In both datasets, the observed values fall outside the null distributions. (E) PCA of the residual matrix (uncorrected minus corrected data) from the CPTAC dataset. Points are colored by laboratory and shaped by biological condition. Residual structure separates by laboratory but not by biological condition, consistent with selective removal of technical variation. (F) Variance decomposition before and after correction. NMFBatch strongly reduces batch-associated variance while retaining sample-associated variation.

Non-negative matrix factorization is a non-convex optimization procedure, and different random initializations may in principle produce different solutions. To address this, we ran NMFBatch 500 times on the CPTAC spike-in dataset using independent random seeds and calculated pairwise sample correlations across runs. The resulting correlations, stratified by laboratory, were tightly clustered, with a mean of 0.99 and a standard deviation of 0.005 as displayed in Fig. 4C, indicating that the NMFBatch-corrected protein profiles were highly reproducible across initializations.

We next tested whether a GAM would overfit patterns by producing biological clustering from random labels (i.e. overfitting). After one NMFBatch correction, we generated 2,000 random permutations by shuffling the condition and laboratory labels. The condition silhouette score was outside the permutation distribution in the CPTAC dataset (permutation P < 0.001; Fig. 4D; left panel). The balanced Quartet dataset gave a similar result (permutation P < 0.001; Fig. 4D; right panel). These analyses provide no evidence that the observed cluster separation was generated randomly e.g. by overfitting.

A PCA of the residual matrix, defined as the uncorrected matrix minus the corrected matrix, showed separation mainly by laboratory rather than by spike-in condition (Fig. 4E). This indicates that the removed component was predominantly associated with interlaboratory variation. PVCA further showed that batch-associated variance decreased from 56.35% to 0.27%, while condition-associated variance increased from 9.21% to 17.38% (Fig. 4F). Pairwise Pearson correlations showed the same overall pattern (Figs. S6 and S7).

Together, these results show that NMFBatch was reproducible across random initializations and reduced laboratory-associated variation while retaining the known concentration-dependent signal in the CPTAC dataset. This indicates that its correction is not attributable to overfitting from the NMF or GAM component.

### NMFBatch supports missing-value imputation and downstream differential-abundance analysis

We first examined whether correction and imputation improved recovery of known spike-in differences in the CPTAC dataset. We performed pairwise differential-abundance testing with limma under four scenarios: NMFBatch correction without imputation, NMFBatch correction with on-the-fly imputation, no matrix correction with laboratory included as a model covariate, and no matrix correction with laboratory included as a covariate after BPCA imputation. PCA showed that NMFBatch correction with imputation reduced separation by laboratory while preserving separation among the spike-in concentration groups (Fig. 5A), consistent with the results in Fig. 4.

**Figure 5.**
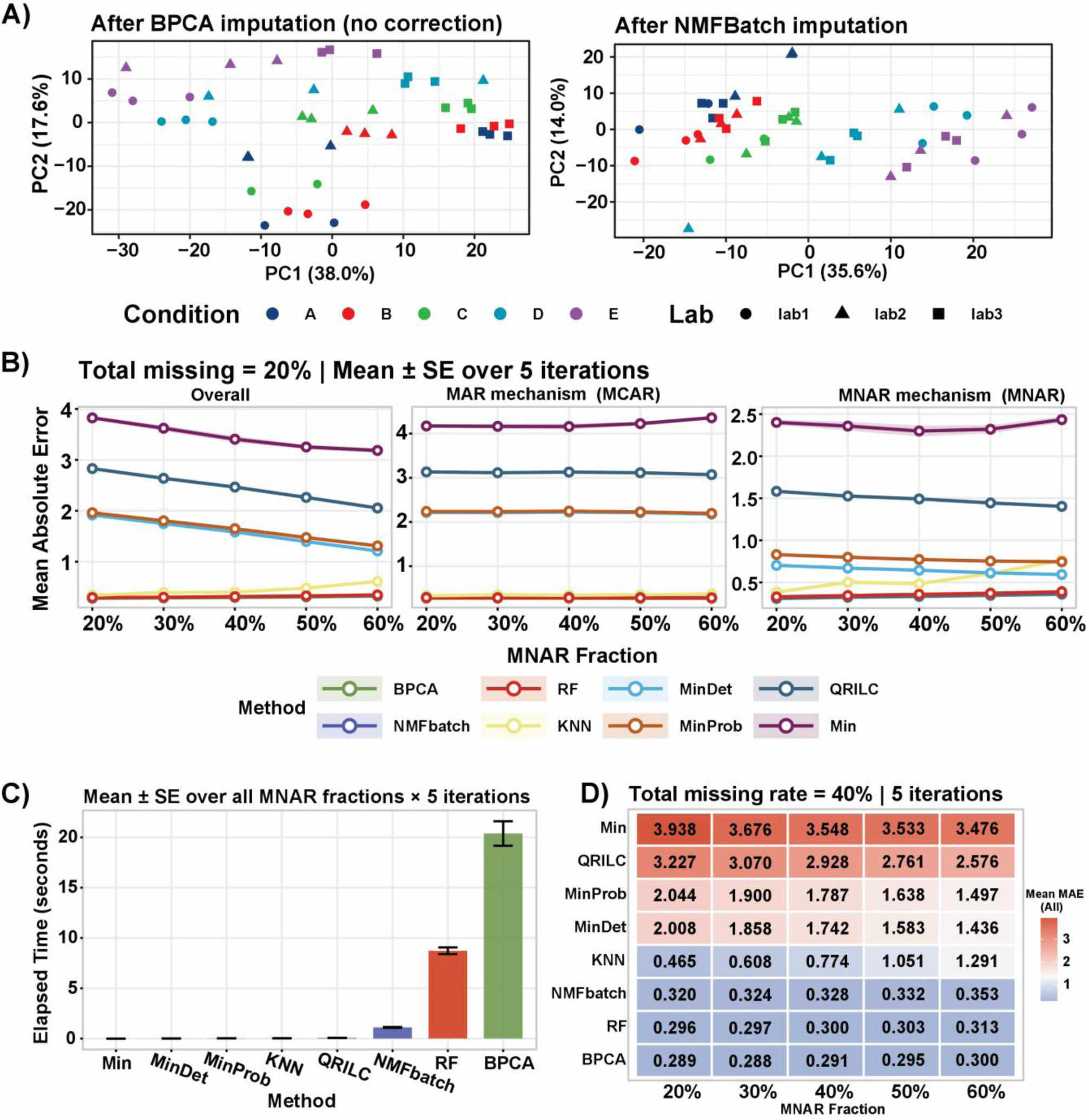
Evaluation of NMFBatch missing-value scoring and imputation performance under simulated MCAR and MNAR missingness. (A) PCA plots showing the sample distribution across spike-in concentration conditions and laboratories. Colors represent the different spike-in concentration conditions, and shapes represent the laboratories of origin. The left panel shows the uncorrected data after BPCA imputation, whereas the right panel shows the data after NMFBatch correction and imputation. NMFBatch reduced laboratory-associated batch effects while preserving the variation associated with the spike-in concentration conditions. (B) Imputation accuracy benchmark across methods under simulated MCAR and MNAR missingness conditions. Mean absolute error (MAE) was calculated between masked true values and imputed values as the proportion of MNAR missingness increased from 20% to 60% while total missingness was fixed at 20%. Methods compared include RF, BPCA, KNN, MinDet, MinProb, QRILC, minimum-value imputation (Min), and NMFBatch on-the-fly imputation. Under MCAR conditions, NMFBatch on-the-fly imputation, RF, and BPCA showed the lowest MAE. Under MNAR conditions. (C) Computational time for the imputation methods shown in panel B. Elapsed time is reported in seconds. NMFBatch on-the-fly imputation required substantially less computation time than RF and BPCA. (D) Imputation accuracy benchmark across methods under simulated MCAR and MNAR missingness conditions. Mean absolute error (MAE) was calculated between masked true values and imputed values as the proportion of MNAR missingness increased from 20% to 60% while total missingness was fixed at 40%.

Without imputation, NMFBatch correction produced an average recall of 0.85 and an average false discovery rate of 0.24. However, only five UPS proteins could be tested on average because proteins containing missing values were excluded. When no matrix correction was applied and laboratory was included in the limma model, recall was 0.83 and the false discovery rate was 0.20, again with limited protein coverage.

With imputation, NMFBatch produced an average recall of 0.80 and an average false discovery rate of 0.21. BPCA imputation without matrix correction produced the same average recall of 0.80 and a false discovery rate of 0.20. Both imputation-based approaches allowed an average of 29 UPS proteins to be tested (Table S; Figs. S8 and S9). Thus, imputation substantially increased the number of testable spike-in proteins, with a modest reduction in recall relative to the non-imputed analyses.

We then compared NMFBatch on-the-fly imputation with random forest (RF), Bayesian PCA (BPCA), K-nearest neighbors (KNN), MinDet, MinProb, QRILC, and minimum-value substitution. We started with complete proteins from the CPTAC protein-level dataset and simulated overall missingness levels of 20% and 40%. Within each overall level, the MNAR proportion ranged from 20% to 60%, with the remaining simulated values treated as MCAR. Each scenario was repeated independently five times.

At 20% overall missingness, NMFBatch on-the-fly imputation, RF, and BPCA produced the lowest mean absolute errors (MAE) for MCAR values (Fig. 5B, middle panel). These methods also showed the lowest or among the lowest errors across the tested MNAR proportions and when all simulated missing values were analyzed together (Fig. 5B, left and right panels). Runtime measurements under the same settings showed that NMFBatch required less time than RF and BPCA (Fig. 5C). At 40% overall missingness, NMFBatch, BPCA, RF, and KNN showed the best overall performance across the tested mixtures of MCAR and MNAR values (Fig. 5D).

These simulations evaluate reconstruction accuracy under the specific masking procedures used here. They do not establish that a single imputation method is optimal for every biological dataset or every missingness mechanism.

### Reduction of noise due to technical variation improves biological structure in single-cell proteomics data

The pSCoPE single-cell dataset provided a test under high sparsity and multiple technical variables. Before correction, both hierarchical clustering and PCA showed structure associated with technical annotations (Fig. 6A and C, top panel). After NMFBatch correction, the heatmap showed less alignment with LC batch, while patterns associated with cell type became clearer (Fig. 6B). PCA showed a similar change: separation by LC batch decreased and clustering by cell type became more apparent (Fig. 6C, bottom panel). PVCA supported these visual observations. Batch-associated variance decreased from 51.38% before correction to 18.8% after correction, while cell-type-associated variance increased from 33.14% to 38.3% (Fig. 6D).

**Figure 6.**
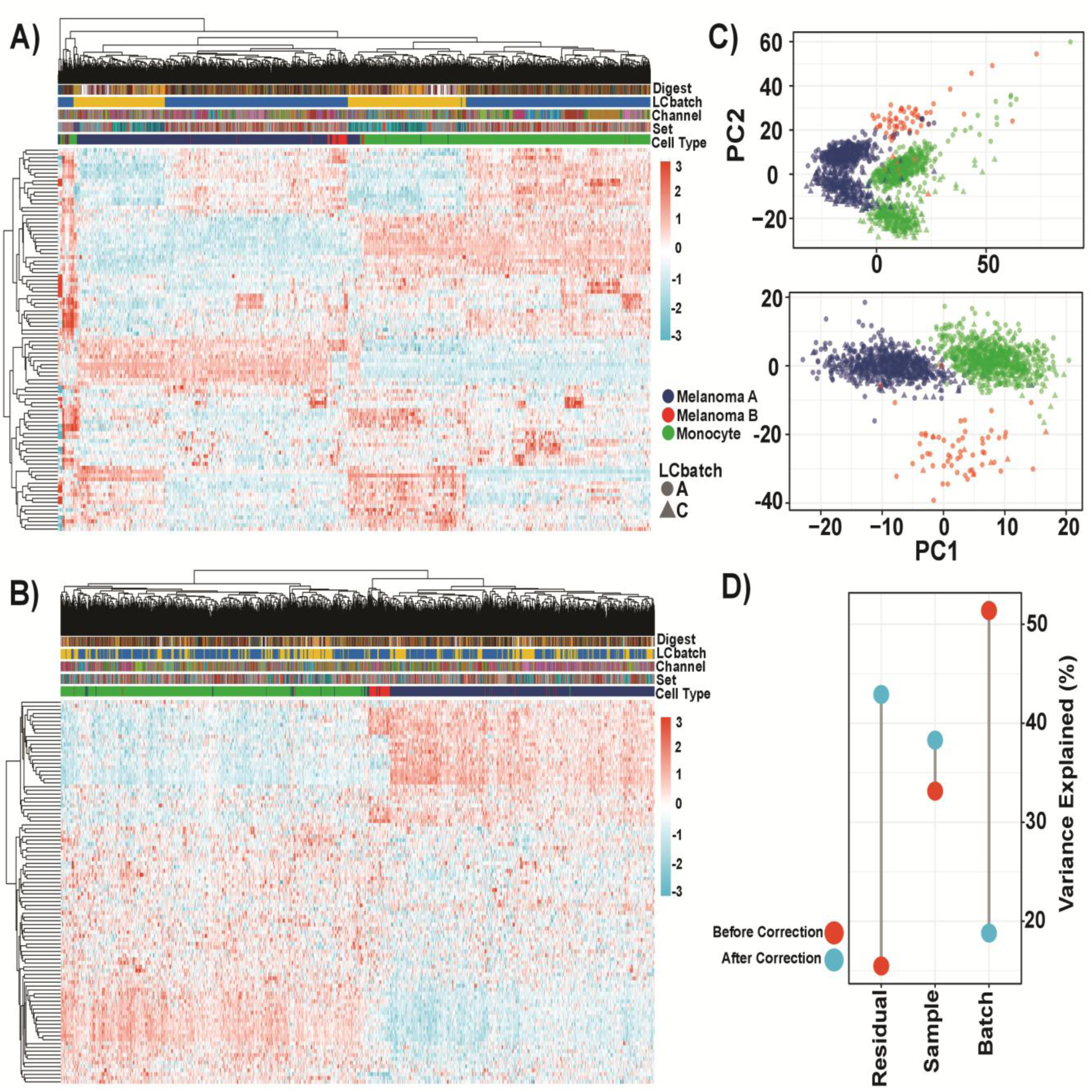
Application of NMFBatch to single-cell proteomics data reduces technical variation while preserving cell-type structure. (A) Hierarchically clustered heatmap of the pSCoPE single-cell proteomics dataset before correction, using top 100 variable proteins. Columns represent single cells and rows represent proteins. Annotation tracks indicate digest, LC batch, TMT channel, labeling set, and cell type. The uncorrected data display prominent structured variation associated with technical covariates. (B) Heatmap of the same dataset after NMFBatch correction, using top 100 variable proteins. Large-scale technical structure is reduced, whereas cell-type-associated patterns remain evident. (C) PCA of the pSCoPE data before correction (top) and after NMFBatch correction (bottom). Samples are coloured by cell type and shaped by LC batch. After correction, batch-associated separation is reduced, and cell-type clustering becomes more apparent. (D) Variance decomposition before and after correction. NMFBatch decreases the batch-associated variance component while preserving sample-associated variance.

These results indicate that NMFBatch reduced, but did not completely eliminate, measured technical variation in this sparse single-cell dataset while preserving and clarifying cell-type-associated structure.

## Discussion

NMFBatch introduces, to our knowledge, the first unified framework that jointly models discrete and continuous sources of unwanted variation, both linear and non-linear, using NMF and GAMs. We illustrate this using large-scale MS proteomics experiments work on protein level data. This protein-level strategy is consistent with a recent benchmarking study supporting protein-level correction for integration of multi-batch proteomics datasets ^8^. NMFBatch masks missing values during model fitting. Consequently, batch effects are estimated from the observed measurements rather than from values imputed before correction. After correction, users can either retain the values reconstructed during NMF or restore the original missing-value pattern and apply another imputation method.

Across the evaluated datasets, NMFBatch reduced technical variation while retaining biological structure. Its largest relative advantage appeared in the confounded Quartet design, in which batch and sample type were correlated. In the balanced design, simpler methods such as median centering also performed well, and the differences between methods were small. In the confounded design, however, several methods removed the measured batch component but retained less sample-associated variation than NMFBatch. NMFBatch also produced the highest median ICC in this setting. These results indicate that removing estimated batch variance alone is not sufficient to establish successful correction because a method must also preserve variation associated with the biological factor of interest.

One possible explanation for the performance of NMFBatch under confounding is that it estimates unwanted variation in a low-dimensional latent space rather than correcting each protein independently. The latent factors represent shared patterns across many proteins providing a structured regularization which helps distinguish between batch and biology. However, latent factorization cannot resolve a design in which batch and biology are completely confounded. The favorable results reported here apply to a partially confounded design and should not be interpreted as evidence that NMFBatch can identify biological and technical effects under conditions where the experimental design is confounded.

The GAM component allows NMFBatch to model discrete batch shifts and continuous injection-order drift within one procedure. In the ChiHOPE dataset, jointly modelling batch and injection batch worked as well as a two-stage workflow using LOESS followed by a separate batch-correction method. The penalized smoothers used by the GAM can shrink the correction of drift towards zero when the data shows limited evidence of drift. This limits over-correction in batches with weak order-dependent variation. In the present analysis, residual drift remained visible after several two-stage LOESS workflows but was not evident after the joint NMFBatch model. This suggests that globally fitted splines (GAM) may, in some settings, capture the underlying curvature of instrument drift more effectively than locally weighted regression (LOESS).

The CPTAC analyses addresses two concerns associated with NMF-based correction: sensitivity to random initialization and overfitting by a flexible model. Across 500 independent initializations, the results remained highly stable (correlation of 0.99 and standard deviation of 0.005). This indicates that the model produces reliable results. In addition, the observed biological silhouette scores were outside the null distributions generated from 2,000 label permutations in both CPTAC and Quartet. These results show that the variation produced after correcting the data with NMFBatch were based on true biological structure and not random fake patterns. They do not, however, rule out every possible form of overfitting at NMF ranks.

The CPTAC spike-in experiment also showed that NMFBatch reduced laboratory-associated variation while retaining the known concentration-dependent signal. Differential-abundance analysis was included as a controlled validation of signal preservation rather than as a general recommendation to perform inference on batch-corrected matrices. Without imputation, recall was slightly higher, but only a small number of UPS proteins could be tested because proteins with missing values were excluded. NMFBatch and BPCA imputation increased the average number of testable UPS proteins from approximately five to 29, with a modest reduction in recall and similar false discovery rates.

In a head-to-head test with simulated data, NMFBatch, RF, and BPCA all performed well, showing low reconstruction errors at 20% overall missingness, although these tools and KNN performed best at 40% missingness. NMFBatch also required less computation time than RF and BPCA under the tested settings. These comparisons describe performance for protein-level values masked according to the simulation procedure used in this study. They should not be interpreted as proving that NMFBatch is optimal for all real-world datasets or missingness mechanisms.

We recognise that imputation at the protein level is conceptually different from imputation at the peptide level. At the peptide level, missingness may reflect low abundance, peptide-specific detectability, interference, or stochastic acquisition. These represent MNAR and MCAR cases. A summarized protein value combines information from multiple peptides. Thus, a missing protein value it is usually due to a complex combination of missingness patterns. Accordingly, the protein-level MCAR and MNAR labels used in our simulations are operational categories for the missing protein measurements, not evidence that every contributing peptide followed the same mechanism.

Several limitations should be considered. First, selecting NMFBatch number of factors relies on user judgement. Although the NMFBatch tool offers diagnostic metrics to guide this selection; different selections can introduce analytical variability. Second, estimating latent factors requires sufficient data. In datasets such as highly sparse single-cell proteomics data, too few observations will lead to poorly constrained factor estimates. The 70% missingness threshold applied to this dataset was a test case to evaluate model stability under sparsity, not a universal cutoff or rule.

The appropriate threshold will depend on the characteristics of the dataset, including sample size, overall sparsity, and the selected NMF factors. This should be determined according to the specific data and model configuration.

Future development of the tool should focus on data-adaptive selection of the missingness threshold and the NMF number of factors. This would ensure that sufficient data is retained to keep the model stable. Furthermore, in future releases, we plan to support NMFBatch through Bioconductor and to maintain the package on both Codeberg and Bioconductor.

## Conclusions

The results we presented here suggest that NMFBatch provides an effective and flexible framework for batch correction in proteomics using protein level data, modelling both batch structure and continuous drift within a single approach. By operating directly on incomplete matrices, supporting on-the-fly missing-value imputation, and maintaining numerical stability across repeated runs, NMFBatch performs competitively under confounded designs while strictly preserving biological variation. These capabilities make it particularly well suited for complex large-scale and single-cell proteomics studies. Although factor selection and performance in extremely sparse datasets remain important practical considerations, these findings position NMFBatch as a promising approach for proteomics studies in which technical heterogeneity and missingness would otherwise limit downstream biological interpretation. More broadly, this framework shows potential as a cross-dataset harmonization framework, enabling integration of proteomics data generated across different cohorts, laboratories, platforms, or acquisition strategies.

## Abbreviations

MS: Mass spectrometry
MCAR: missing completely at random
MAR: missing at random
MNAR: missing not at random
LOESS: Locally Estimated Scatterplot Smoothing
NMF: non-negative matrix factorization
GAMs: generalized additive models
ANLS: non-negative least squares
NNLS: non-negative least squares
SCD: sequential coordinate-wise descent
REML: restricted maximum likelihood
PVCA: Principal Variance Component Analysis
ICC: Intraclass Correlation Coefficient
PCA: Principal Component Analysis
DDA: data-dependent acquisition
DIA: data-independent acquisition
CPTAC: Clinical Proteomic Technology Assessment for Cancer

## Declarations

### Ethics approval and consent to participate

This study used publicly available proteomics datasets and performed secondary analyses only. Ethical approval and consent were addressed in the original studies.

### Consent for publication

Not applicable

### Availability of data and materials

All data used in this study are publicly accessible. The Quartet uncorrected and batch-corrected datasets ^8^ are hosted on Figshare https://doi.org/10.6084/m9.figshare.29567333.v2, and the ChiHOPE uncorrected and batch-corrected datasets ^8^ are available at https://doi.org/10.6084/m9.figshare.30028336. The CPTAC dataset ^17^ was obtained from https://github.com/statOmics/GMFProteomicsPaper/tree/main/Data/CPTAC_data ^28^ and is additionally available through the MsDataHub package. The Leduc pSCoPE single-cell proteomics dataset ^4^ was retrieved via the scpdata package. Finally, R scripts are available in the Codeberg repository https://codeberg.org/AliYoussef/NMFBatchManuscript to facilitate reproducibility of the analyses.

### Code availability

NMFBatch is released as an open-source R package and can be accessed at Codeberg https://codeberg.org/AliYoussef/NMFBatch. The accompanying repository https://codeberg.org/AliYoussef/NMFBatchManuscript contains all necessary R scripts and data to reproduce the analyses and results presented in this study. To facilitate reproducibility, we also deposited all datasets and the code required to run the analyses presented in this study in Zenodo: https://doi.org/10.5281/zenodo.20282106.

### Authors’ contributions

Ali Mostafa Anwar (Conceptualization [lead], Formal analysis [lead], Methodology [lead], Software [lead], Visualization [lead], Writing—original draft [lead]), Salma Bayoumi (Software [lead], Validation [supporting], Writing—review & editing [supporting]), Leo Lahti (Supervision [lead], Software [supporting], Validation [supporting], Writing—review & editing [supporting]), and Eleanor Coffey (Supervision [lead], Investigation [supporting], Validation [supporting], Writing—review & editing [supporting]). All authors read and approved the final manuscript.

## Acknowledgement

GPT-5.4 was used for English language and grammar correction. Claude Opus 4.6 assisted only with writing NMFBatch package documentation during development. All documentation generated with AI was reviewed by A.M.A and S.B. The NMFBatch logo used in the graphical abstract was generated using Nano Banana 2, Gemini’s AI image generator, and edited in CorelDRAW.

## Supplementary information

Tables from S1 to S5 and Supplementary Figures S1-S14.

## References

(1) Williams, E. G.; Pfister, N.; Roy, S.; Statzer, C.; Haverty, J.; Ingels, J.; Bohl, C.; Hasan, M.; Čuklina, J.; Bühlmann, P.; Zamboni, N.; Lu, L.; Ewald, C. Y.; Williams, R. W.; Aebersold, R. Multiomic Profiling of the Liver across Diets and Age in a Diverse Mouse Population. Cell Syst. 2022, 13 (1), 43–57.e6. 10.1016/J.CELS.2021.09.005

(2) He, F.; Aebersold, R.; Baker, M. S.; Bian, X.; Bo, X.; Chan, D. W.; Chang, C.; Chen, L.; Chen, X.; Chen, Y.-J.; Cheng, H.; Collins, B. C.; Corrales, F.; Cox, J.; E, W.; Van Eyk, J. E.; Fan, J.; Faridi, P.; Figeys, D.; Gao, G. F.; Gao, W.; Gao, Z.-H.; Goda, K.; Goh, W. W. Bin; Gu, D.; Guo, C.; Guo, T.; He, Y.; Heck, A. J. R.; Hermjakob, H.; Hunter, T.; Iyer, N. G.; Jiang, Y.; Jimenez, C. R.; Joshi, L.; Kelleher, N. L.; Li, M.; Li, Y.; Lin, Q.; Liu, C. H.; Liu, F.; Liu, G.-H.; Liu, Y.; Liu, Z.; Low, T. Y.; Lu, B.; Mann, M.; Meng, A.; Moritz, R. L.; Nice, E.; Ning, G.; Omenn, G. S.; Overall, C. M.; Palmisano, G.; Peng, Y.; Pineau, C.; Poon, T. C. W.; Purcell, A. W.; Qiao, J.; Reddel, R. R.; Robinson, P. J.; Roncada, P.; Sander, C.; Sha, J.; Song, E.; Srivastava, S.; Sun, A.; Sze, S. K.; Tang, C.; Tang, L.; Tian, R.; Vizcaíno, J. A.; Wang, C.; Wang, C.; Wang, X.; Wang, X.; Wang, Y.; Weiss, T.; Wilhelm, M.; Winkler, R.; Wollscheid, B.; Wong, L.; Xie, L.; Xie, W.; Xu, T.; Xu, T.; Yan, L.; Yang, J.; Yang, X.; Yates, J.; Yun, T.; Zhai, Q.; Zhang, B.; Zhang, H.; Zhang, L.; Zhang, L.; Zhang, P.; Zhang, Y.; Zheng, Y. Z.; Zhong, Q.; Zhu, Y.; Chan, D. W.; Tan, C. S. H.; E, W.; Van Eyk, J. E.; Gao, M.; Gao, Q.; Gao, Y.; Guo, X.; He, J.; He, J.; He, Q.-Y.; Hou, J.; Huang, C.; Jia, C.; Kuster, B.; Li, C.; Li, D.; Li, Y.; Li, Y.; Liu, S.; Liu, X.; Liu, Y.; Liu, Z.; Lu, H.; Nie, Y.; Okada, M.; Qian, G.; Qin, H.; Rao, Y.; Rao, Z.; Ren, X.; Ren, Y.; Shen, F.; Shen, L.; Su, S.; Tan, M.; Tang, B. Z.; Tao, S.-C.; Wang, J.; Wang, T.; Wei, L.; Wong, C. C. L.; Xie, X. S.; Xu, L.; Xu, P.; Xu, T.; Yang, H.; Yang, J.; Ye, M.; Ying, W.; Yu, X.; Yuan, Y.; Zeng, Q.; Zhan, Q.; Zhang, X.; Zhang, X.; Zhang, Y.; Zhong, N.-S.; Zhou, F.; Zhu, Y.; Zhang, Z. Author Correction: π-HuB: The Proteomic Navigator of the Human Body. Nature 2024 637:8046 2024, 637 (8046), E22–E22. 10.1038/s41586-024-08555-x

(3) Liu, Y.; Buil, A.; Collins, B. C.; Gillet, L. C.; Blum, L. C.; Cheng, L.; Vitek, O.; Mouritsen, J.; Lachance, G.; Spector, T. D.; Dermitzakis, E. T.; Aebersold, R. Quantitative Variability of 342 Plasma Proteins in a Human Twin Population. Mol. Syst. Biol. 2015, 11 (2), MSB145728-. 10.15252/MSB.20145728/FIGURES/5

(4) Leduc, A.; Huffman, R. G.; Cantlon, J.; Khan, S.; Slavov, N. Exploring Functional Protein Covariation across Single Cells Using NPOP. Genome Biology 2022 23:1 2022, 23 (1), 261-. 10.1186/S13059-022-02817-5

(5) Meissner, F.; Geddes-McAlister, J.; Mann, M.; Bantscheff, M. The Emerging Role of Mass Spectrometry-Based Proteomics in Drug Discovery. Nat. Rev. Drug Discov. 2022, 21 (9), 637–654. 10.1038/s41573-022-00409-3

(6) Poulos, R. C.; Hains, P. G.; Shah, R.; Lucas, N.; Xavier, D.; Manda, S. S.; Anees, A.; Koh, J. M. S.; Mahboob, S.; Wittman, M.; Williams, S. G.; Sykes, E. K.; Hecker, M.; Dausmann, M.; Wouters, M. A.; Ashman, K.; Yang, J.; Wild, P. J.; deFazio, A.; Balleine, R. L.; Tully, B.; Aebersold, R.; Speed, T. P.; Liu, Y.; Reddel, R. R.; Robinson, P. J.; Zhong, Q. Strategies to Enable Large-Scale Proteomics for Reproducible Research. Nature Communications 2020 11:1 2020, 11 (1), 3793-. 10.1038/s41467-020-17641-3

7. (7) Goh, W. W. Bin; Wang, W.; Wong, L. Why Batch Effects Matter in Omics Data, and How to Avoid Them. Trends Biotechnol. 2017, 35 (6), 498–507. 10.1016/J.TIBTECH.2017.02.012

(8) Chen, Q.; Cao, Z.; Liu, Y.; Zhang, N.; Xie, Y.; Chen, H.; Mai, Y.; Duan, S.; Li, J.; Yu, Y.; Zhao, Y.; Shi, L.; Zheng, Y. Protein-Level Batch-Effect Correction Enhances Robustness in MS-Based Proteomics. Nature Communications 2025 16:1 2025, 16 (1), 9735-. 10.1038/s41467-025-64718-y

(9) Čuklina, J.; Lee, C. H.; Williams, E. G.; Sajic, T.; Collins, B. C.; Rodríguez Martínez, M.; Sharma, V. S.; Wendt, F.; Goetze, S.; Keele, G. R.; Wollscheid, B.; Aebersold, R.; Pedrioli, P. G. A. Diagnostics and Correction of Batch Effects in Large-scale Proteomic Studies: A Tutorial. Molecular Systems Biology 2021 17:8 2021, 17 (8), MSB202110240-. 10.15252/MSB.202110240

(10) Karpievitch, Y. V.; Dabney, A. R.; Smith, R. D. Normalization and Missing Value Imputation for Label-Free LC-MS Analysis. BMC Bioinformatics 2012 13:16 2012, 13 (16), S5-. 10.1186/1471-2105-13-S16-S5

(11) Matafora, V.; Corno, A.; Ciliberto, A.; Bachi, A. Missing Value Monitoring Enhances the Robustness in Proteomics Quantitation. J. Proteome Res. 2017, 16 (4), 1719–1727. 10.1021/ACS.JPROTEOME.6B01056

(12) Johnson, W. E.; Li, C.; Rabinovic, A. Adjusting Batch Effects in Microarray Expression Data Using Empirical Bayes Methods. Biostatistics 2007, 8 (1), 118–127. 10.1093/BIOSTATISTICS/KXJ037

(13) Voß, H.; Schlumbohm, S.; Barwikowski, P.; Wurlitzer, M.; Dottermusch, M.; Neumann, P.; Schlüter, H.; Neumann, J. E.; Krisp, C. HarmonizR Enables Data Harmonization across Independent Proteomic Datasets with Appropriate Handling of Missing Values. Nature Communications 2022 13:1 2022, 13 (1), 3523-. 10.1038/s41467-022-31007-x

(14) Ritchie, M. E.; Phipson, B.; Wu, D.; Hu, Y.; Law, C. W.; Shi, W.; Smyth, G. K. Limma Powers Differential Expression Analyses for RNA-Sequencing and Microarray Studies. Nucleic Acids Res. 2015, 43 (7), e47–e47. 10.1093/nar/gkv007

(15) Zheng, Y.; Liu, Y.; Yang, J.; Dong, L.; Zhang, R.; Tian, S.; Yu, Y.; Ren, L.; Hou, W.; Zhu, F.; Mai, Y.; Han, J.; Zhang, L.; Jiang, H.; Lin, L.; Lou, J.; Li, R.; Lin, J.; Liu, H.; Kong, Z.; Wang, D.; Dai, F.; Bao, D.; Cao, Z.; Chen, Q.; Chen, Q.; Chen, X.; Gao, Y.; Jiang, H.; Li, B.; Li, B.; Li, J.; Liu, R.; Qing, T.; Shang, E.; Shang, J.; Sun, S.; Wang, H.; Wang, X.; Zhang, N.; Zhang, P.; Zhang, R.; Zhu, S.; Scherer, A.; Wang, J.; Wang, J.; Huo, Y.; Liu, G.; Cao, C.; Shao, L.; Xu, J.; Hong, H.; Xiao, W.; Liang, X.; Lu, D.; Jin, L.; Tong, W.; Ding, C.; Li, J.; Fang, X.; Shi, L. Multi-Omics Data Integration Using Ratio-Based Quantitative Profiling with Quartet Reference Materials. Nature Biotechnology 2023 42:7 2023, 42 (7), 1133–1149. 10.1038/s41587-023-01934-1

(16) Yang, J.; Liu, Y.; Shang, J.; Chen, Q.; Chen, Q.; Ren, L.; Zhang, N.; Yu, Y.; Li, Z.; Song, Y.; Yang, S.; Scherer, A.; Tong, W.; Hong, H.; Xiao, W.; Shi, L.; Zheng, Y. The Quartet Data Portal: Integration of Community-Wide Resources for Multiomics Quality Control. Genome Biology 2023 24:1 2023, 24 (1), 245-. 10.1186/S13059-023-03091-9

(17) Paulovich, A. G.; Billheimer, D.; Ham, A. J. L.; Vega-Montoto, L.; Rudnick, P. A.; Tabb, D. L.; Wang, P.; Blackman, R. K.; Bunk, D. M.; Cardasis, H. L.; Clauser, K. R.; Kinsinger, C. R.; Schilling, B.; Tegeler, T. J.; Variyath, A. M.; Wang, M.; Whiteaker, J. R.; Zimmerman, L. J.; Fenyo, D.; Carr, S. A.; Fisher, S. J.; Gibson, B. W.; Mesri, M.; Neubert, T. A.; Regnier, F. E.; Rodriguez, H.; Spiegelman, C.; Stein, S. E.; Tempst, P.; Liebler, D. C. Interlaboratory Study Characterizing a Yeast Performance Standard for Benchmarking LC-MS Platform Performance. Molecular and Cellular Proteomics 2010, 9 (2), 242–254. 10.1074/mcp.M900222-MCP200

(18) Lee, D. D.; Seung, H. S. Learning the Parts of Objects by Non-Negative Matrix Factorization. Nature 1999 401:6755 1999, 401 (6755), 788–791. 10.1038/44565

(19) Brunet, J. P.; Tamayo, P.; Golub, T. R.; Mesirov, J. P. Metagenes and Molecular Pattern Discovery Using Matrix Factorization. Proceedings of the National Academy of Sciences 2004, 101 (12), 4164–4169. 10.1073/PNAS.0308531101

(20) Kim, H.; Park, H. Sparse Non-Negative Matrix Factorizations via Alternating Non-Negativity-Constrained Least Squares for Microarray Data Analysis. Bioinformatics 2007, 23 (12), 1495–1502. 10.1093/BIOINFORMATICS/BTM134

(21) Alexandrov, L. B.; Nik-Zainal, S.; Wedge, D. C.; Campbell, P. J.; Stratton, M. R. Deciphering Signatures of Mutational Processes Operative in Human Cancer. Cell Rep. 2013, 3 (1), 246–259. 10.1016/J.CELREP.2012.12.008

(22) Lin, X.; Boutros, P. C. Optimization and Expansion of Non-Negative Matrix Factorization. BMC Bioinformatics 2019 21:1 2020, 21 (1), 7-. 10.1186/S12859-019-3312-5

(23) Lazar, C.; Burger, T. A Collection of Methods for Left-Censored Missing Data Imputation [R Package ImputeLCMD Version 2.1]. CRAN: Contributed Packages 2022. 10.32614/CRAN.PACKAGE.IMPUTELCMD

(24) Huber, W.; Carey, V. J.; Gentleman, R.; Anders, S.; Carlson, M.; Carvalho, B. S.; Bravo, H. C.; Davis, S.; Gatto, L.; Girke, T.; Gottardo, R.; Hahne, F.; Hansen, K. D.; Irizarry, R. A.; Lawrence, M.; Love, M. I.; MaCdonald, J.; Obenchain, V.; Oles, A. K.; Pagès, H.; Reyes, A.; Shannon, P.; Smyth, G. K.; Tenenbaum, D.; Waldron, L.; Morgan, M. Orchestrating High-Throughput Genomic Analysis with Bioconductor. Nature Methods 2015 12:2 2015, 12 (2), 115–121. 10.1038/nmeth.3252

(25) Chen, Q. All Corrected/Uncorrected Data Matrices at Precursor, Peptide, Protein Levels. 2025. 10.6084/m9.figshare.29567333.v2

(26) Chen, Q. All Preprocessed ChiHOPE Data Matrices. 2025. 10.6084/m9.figshare.30028336.v1

(27) Demichev, V.; Messner, C. B.; Vernardis, S. I.; Lilley, K. S.; Ralser, M. DIA-NN: Neural Networks and Interference Correction Enable Deep Proteome Coverage in High Throughput. Nature Methods 2019 17:1 2019, 17 (1), 41–44. 10.1038/s41592-019-0638-x

(28) Segers, A.; Castiglione, C.; Vanderaa, C.; Martens, L.; Risso, D.; Clement, L. OmicsGMF: A Multi-Tool for Dimensionality Reduction, Batch Correction and Imputation Applied to Bulk-and Single Cell Proteomics Data. bioRxiv 2025, 2025.03.24.644996. 10.1101/2025.03.24.644996

(29) Vanderaa, C.; Gatto, L. The Current State of Single-Cell Proteomics Data Analysis. Curr. Protoc. 2023, 3 (1), e658. 10.1002/CPZ1.658

(30) Korsunsky, I.; Millard, N.; Fan, J.; Slowikowski, K.; Zhang, F.; Wei, K.; Baglaenko, Y.; Brenner, M.; Loh, P. ru; Raychaudhuri, S. Fast, Sensitive and Accurate Integration of Single-Cell Data with Harmony. Nature Methods 2019 16:12 2019, 16 (12), 1289–1296. 10.1038/s41592-019-0619-0

(31) Deng, K.; Zhao, F.; Rong, Z.; Cao, L.; Zhang, L.; Li, K.; Hou, Y.; Zhu, Z. J. WaveICA 2.0: A Novel Batch Effect Removal Method for Untargeted Metabolomics Data without Using Batch Information. Metabolomics 2021 17:10 2021, 17 (10), 87-. 10.1007/S11306-021-01839-7

